# Parallel and non-parallel divergence within polymorphic populations of brook stickleback, *Culaea inconstans* (Actinopterygii: Gasterosteidae)

**DOI:** 10.1101/2021.06.08.447580

**Authors:** Kaitlyn Willerth, Emily Franks, Jonathan A. Mee

## Abstract

Studying parallel evolution allows us to draw conclusions about the repeatability of adaptive evolution. Whereas populations likely experience similar selective pressures in similar environments, it is not clear if this will always result in parallel divergence of ecologically relevant traits. Our study investigates the extent of parallelism associated with the evolution of pelvic spine reduction in brook stickleback populations. We find that populations with parallel divergence in pelvic spine morphology do not exhibit parallel divergence in head and body morphology but do exhibit parallel divergence in diet. In addition, we compare these patterns associated with pelvic reduction in brook stickleback to well-studied patterns of divergence between spined and unspined threespine stickleback. Whereas spine reduction is associated with littoral habitats and a benthic diet in threespine stickleback, spine reduction in brook stickleback is associated with a planktonic diet. Hence, we find that pelvic spine divergence is associated with largely non-parallel ecological consequences across species.

## Introduction

Understanding the causes of adaptive phenotypic divergence is a central goal in evolutionary biology. The genetic basis of adaptive divergence has received much needed attention recently, but the ecological causes of selection on divergent phenotypes remain unknown in many cases. Whereas understanding the genetic basis of phenotypic variation is central to many questions in evolutionary biology, understanding the ecological correlates of adaptive variation is key to answering questions about the causes of selection that drive diversification. In populations with persistent adaptive phenotypic dimorphism, selection may involve several ecological factors simultaneously, including predation, competition, or parasitism, and selection driving divergence in one trait may affect divergence in other traits due to pleiotropy. Alternatively, a suite of traits may change in concert with a particular adaptive polymorphism due to plastic or evolutionary responses in resource use, habitat use, behaviour, or predator interactions that arise as a consequence of the trait polymorphism. For example, cliff nesting in some kittiwake gulls (*Rissa tridactyla* Linnaeus 1758) results in reduced predation risk and reduced predator avoidance behaviours (Cullen 1956). The diurnal life history of butterflies likely originated as a strategy to avoid bat predation and is associated with the loss of ultrasonic hearing (Yack and Fullard 2008). In threespine stickleback (*Gasterosteus acculeatus* Linnaeus 1758), predators induce selection for longer protective spines, which are part of a suite of traits associated with habitat divergence within some freshwater lakes (Rennison et al. 2019). In all of these examples, predator-related differences in phenotype are associated with other ecological or behavioural differences. Sticklebacks, in particular, have emerged as a model system in the study of adaptive divergence, particularly with respect to the role of predation in driving adaptive divergence.

Three species in the stickleback family (Gasterosteidae), in three different genera (*Gasterosteus, Pungitius*, and *Culaea*), exhibit heritable variation in their pelvic or dorsal spines (Nelson 1969, Nelson 1977, Chan *et al*. 2010). There is good evidence that spines are involved, at least to some extent, in defence against predators (Hoogland et al. 1957, Hall 1956, Reisman and Cade 1967). For example, gape-limited predators (e.g. some birds, trout, and other small or medium-sized fishes) are deterred by spines, and these predators select for increased armor and longer spines in stickleback (Reist 1980a, Reimchen 1992, Vamosi and Schluter 2004, Marchinko 2008, Lescak and von Hippel 2011, Miller et al. 2017). Spined stickleback are bolder and will tolerate being closer to predators than unspined stickleback (Reist 1980*a*, Reist 1980*b*, Reist 1983). Large fish, such as pike (*Esox lucius* Linnaeus 1758), which are not gape-limited, are less deterred by armor and spines. Hence, predators that are not gape-limited likely select for spine reduction or spine loss (Nelson and Atton 1971, Andraso and Barron 1995, Leinonen et al. 2011). Anecdotal evidence has suggested that invertebrate predators, such as dragonfly nymphs (Odonata: Anisoptera) and giant water bugs (Hemiptera: Belostomatidae), use spines and other external bony structures to grasp their prey, suggesting that invertebrates may select for spine reduction or loss (Reimchen 1980, Reist 1980*b*, Vamosi 2002, Lescak et al. 2012), but a meta-analysis of invertebrate selection experiments, found little support for selection against spines (Miller et al. 2017). Several other predators are known to prey on stickleback (Reimchen 1994), including loons (*Gavia immer* Brunnich 1764; Reimchen 1980), muskrats (*Ondatra zibethicus* Linnaeus 1766; Nelson 1977), and conspecific stickleback (Foster et al. 1988), but the influence of each of these predators on stickleback spines is difficult to predict and has not been evaluated. The evidence that predation influences selection on pelvic phenotypes in stickleback is further supported by the observation that the size of pelvic girdle and pelvic spines in stickleback is proportional to the density of predatory fish in a region (Miller *et al*. 2017). Insofar as different predators use different habitats within lakes (Reimchen 1994), predator-mediated selection may drive spined and unspined stickleback into different habitats.

Pelvic spines have received much more attention than dorsal spines in the stickleback literature. Ancestrally, all species of stickleback had a pelvic structure composed of a pelvic girdle and two spines (Bell 1974, Ward and McLennan 2009). Loss of the pelvic spines and pelvic girdle has evolved in hundreds of populations, and many individuals have intermediate or ‘vestigial’ pelvic structures such as half a girdle with only one spine or a complete pelvic girdle with no spines (Klepaker 2013). Divergence in pelvic phenotype within stickleback populations may be associated with resource competition leading to different habitat use (Nelson 1977, Schulter 1994), but balancing selection caused by different predators that select for different traits has been implicated as the main driver of pelvic divergence within and among stickleback populations (Nelson 1969, Nelson and Atton 1971, Reimchen 1980, Reist 1980a, Reist 1980b, Reimchen 1994, Marchinko 2009). In the absence of predators, pelvic reduction is associated with low-calcium environments (Bell et al. 1993), but it is unlikely that variation in calcium availability among habitats within a population would be sufficient to drive within-population spine polymorphism. Populations with pelvic reduction are more often found in lakes which lack at outlet (Nelson and Atton 1971), suggesting, perhaps, that the presence of multiple habitats within are lake are required for pelvic spine divergence. Regardless of whether predation was the initial, or is the primary, cause of the pelvic phenotype divergence within stickleback populations, predators may drive different pelvic phenotypes to use different habitats within a population, and individuals that use different habitats are likely exposed to different environmental effects (see, for example, Rennison et al. 2019). The maintenance of within-population divergence may, therefore, be dependent on multiple ecological factors, and adaptive phenotypic divergence in pelvic morphology may have consequences on a variety of ecologically relevant traits.

We studied populations of brook stickleback (*Culaea inconstans* Kirtland, 1840) to investigate the hypothesis that divergent pelvic phenotypes are associated with divergence in habitat. Pelvic spine polymorphism in threespine stickleback (*Gasterosteus acculeatus* Linnaeus, 1958) has been studied extensively, and, in a few well-studied cases, threespine stickleback with dimorphic pelvic phenotypes are reproductively isolated sympatric ecomorphs (McPhail 1984, Ridgeway and McPhail 1984, McPhail 1992, Schluter and McPhail 1992, Nagel and Schluter 1998, Rundle et al. 2000). In the majority of polymorphic threespine stickleback populations, however, vestigial pelvic phenotypes are more abundant than either the fully spined morph or the unspined morph (Klepaker et al. 2013). In contrast, the vestigial morphs are absent or rare in most polymorphic brook stickleback populations (Nelson and Atton 1971, Nelson 1977, Klepaker et al. 2013). Also, unlike dimorphic threespine stickleback populations, pelvic phenotypes in brook stickleback populations are not reproductively isolated, but, nonetheless, have persisted over multiple generations at stable frequencies except where anthropogenic environmental disturbances have occurred (Lowey et al. 2020). Dorsal and pelvic spines in brook stickleback are longest in the southern parts of their distribution and shortest in the north (Nelson 1969), whereas clinal variation in spine length is not present in threepine. An additional notable difference between brook stickleback and other stickleback species with respect to spine reduction is that there are no marine populations of brook stickleback, and, therefore, spine reduction in brook stickleback is not associated with freshwater colonization (Nelson 1969)

We assessed body shape variation associated with pelvic spine polymorphism in brook stickleback because organisms’ body shapes can be substantially influenced by being exposed to different environments or habitats, and variation in body shape can reflect important ecological and behavioural differences among individuals (Bell and Foster 1994, Reimchen et al. 1985, Webster 2011). In addition, it is possible that the gene or genes involved in pelvic polymorphism may have pleiotropic effects on other morphological traits. We also investigated habitat divergence among brook stickleback pelvic phenotypes by analyzing stable isotope signatures. If the divergent pelvic morphs of brook stickleback use different habitats, then diet is likely to differ as a consequence. According to Cutting *et al* (2016), fish muscle biochemistry reflects a long-term average diet (over a few months) and is a good indicator of diet source. Analysis of carbon (δ13C) and nitrogen (δ15N) isotopes in fish tissues is a common method to evaluate variation in habitat and resource use in freshwater environments (Post 2002). Different photosynthetic organisms (e.g. plants vs phytoplankton) fix carbon isotopes in different ratios. For example, terrestrial plants tend to fix more 13C and, as a result, have higher δ13C values than phytoplankton. When photosynthetic organisms are consumed, their carbon isotope ratios are assimilated and reflected within consumers’ tissues, and δ15N values tend to increase with trophic level due to preferential assimilation of 15N (Jardine et al. 2003; Eloranta et al. 2010). If phenotypically different brook stickleback forage in different habitats (e.g. limnetic vs. benthic), they may have different isotopic signatures if primary producer composition is unique to either habitat and if their diet shifts to higher-order consumers. Based on patterns of ecological divergence between spined and unspined threespine stickleback (Reimchen 1980, McPhail 1992), we predict that brook stickleback with reduced pelvises will be associated with benthic or littoral habitats and will have a more benthic diet, which would lead to higher δ13C values and higher δ15N values among unspined individuals. We do not, however, have a specific prediction about how body shape may change between spined and unspined brook stickleback morphs that use different habitats because, unlike diet and stable isotope signatures, morphological differences between stickeback in different habitats show little parallelism among lakes (Kaeuffer et al. 2012).

## Methods

### Sample preparation and collection

We collected adult brook stickleback from two lakes in Alberta, Canada, in 2017 and 2019 (with UTF-8 encoded WGS84 latitude and longitude): Muir Lake (53.627659, - 113.957524) and Shunda Lake (52.453899, −116.146192). Shunda Lake was previously known as Fish Lake (as in Nelson and Atton 1971, Nelson 1977). These lakes were selected because, among the lakes with polymorphic brook stickleback populations in the region, they had a relatively high abundance of spined and unspined pelvic morphs (Nelson 1977, Lowey et al. 2020). Both lakes have fish survey and stocking records indicating the potential presence of several stickleback predators, including brown trout (*Salmo trutta* Linnaeus 1758), rainbow trout [*Ocorhychus mykiss* (Walbaum 1792)], brook trout (*Salvelinus fontinalis* Mitchill 1814), northern pike (*Esox lucius* Linnaeus 1758), and yellow perch [*Perca flavescens* (Mitchill, 1814)], although recent surveys and stocking records (i.e. since 1990) list only rainbow trout and brown trout, suggesting that these salmonids are likely the dominant predatory fish (Alberta Environment and Parks 2021). In Muir Lake, brook stickleback also coexist with fathead minnow [*Pimephales promelas* (Rafinesque, 1820)], whereas in Shunda Lake the fish community includes longnose sucker (*Catostomus catostomus* Forster 1773), white sucker [*Catostomus commersonii* (Lacépède 1803)], and northern pearl dace [*Margariscus nachtriebi* (Cox 1896)]. We observed loons [*Gavia immer* (Brunnich 1764)], dragonfly nymphs (Gomphidae and other unidentified families), giant water bugs [*Lethocerus americanus* (Leidy 1847)], and backswimmers (Notonectidae) at both lakes. The distributions and abundances of stickleback predators and competitors across habitats in these lakes has not been evaluated. Muir lake has a maximum depth of 6.5m, water conductivity of 236µS/cm, and pH of 8.7 (measured in the summer with water temperature of 18 degrees C), and, although undeveloped native woodlands and residential development surround the lake, the predominant land use in the area is agriculture (Alberta Environment and Parks 2021). Shunda Lake has a maximum depth of 6.2m, water conductivity of 264.5µS/cm, and pH of 8.6 (measured in the summer with water temperature of 16 degrees C), and is surrounded entirely by native woodland (Alberta Environment and Parks 2021). Shunda Lake has an outlet stream, whereas Muir Lake does not.

Brook stickleback were collected in June and July using unbaited minnow traps (5 mm mesh). To sample brook stickleback from the littoral zone, traps were set adjacent to the shore at 0.5-2 m depths. To sample brook stickleback from the limnetic zone, traps were set at least 50 m from shore and suspended from floats at a depth of 1-2m. Traps were retrieved one to twelve hours after being set. All brook stickleback samples were anesthetized and euthanized in an overdose mixture of lake water and eugenol. In 2019, the posterior portion (posterior to the pelvic girdle) of each individual from Muir Lake and Shunda Lake was frozen on dry ice then preserved at −20°C for stable isotope analysis. All other tissues were preserved in 70% EtOH for morphometric analysis. Samples were collected under a fisheries research license issued by the Government of Alberta. Collection methods and the use of animals in research was approved by the Animal Care Committee at Mount Royal University (Animal Care Protocol ID 101029 and 101795). Spined and unspined individuals were initially identified at the site of capture based on close visual inspection and prodding with fine-tipped tweezers. Sex was also assigned at the site of capture by examining gonads and by noting the presence of nuptial colouration. In 2019, benthic invertebrates were collected from the littoral zones of Muir Lake and Shunda Lake by rinsing and sorting through mud samples, and plankton was collected from the pelagic zone of these two lakes using a Wisconsin Plankton Sampler. Benthic invertebrates were identified to family or species (if possible) immediately after capture. Plankton and benthic invertebrate samples were frozen on dry ice immediately, then preserved at −20°C for stable isotope analysis.

### Geometric Morphometrics

The brook stickleback specimens were bleached and dyed using alizarin red following the protocol from Xie *et al*. (2019). After bleaching and staining, we captured ventral and left-lateral photographs of each fish against a 1×1cm grid using a Canon EOS Rebel T6i mounted above each specimen at a height of 15 cm with two SV SlimPanel LED high-intensity illuminators. We verified the pelvic phenotype for each individual by examining the ventral photograph, and, following Kepaker et al. (2013), each individual was classified as having a “normal pelvis” with a complete pelvic girdle and both spines (hereafter “spined”), a “lost pelvis” with complete absence of pelvic girdle and spines (hereafter “absent”), or a “vestigial pelvis” wherein one or more spines or pelvic girdle elements (the ascending branch, anterior process, or posterior process) is missing. The population from Muir Lake contains very few vestigial pelvic phenotypes (2 to 3% of the population), and, to allow comparisons across populations, we combined the absent and vestigial pelvic phenotypes into a single category that we called “reduced”. Analyses involving the three pelvic phenotypes (i.e., spined, vestigial, and absent) were not possible in all instances (i.e. due to the lack of vestigial individuals in Muir Lake samples), and, when they were possible, did not yield substantial differences in statistical results or overall conclusions relative to the analyses with two categories (i.e. spined and reduced) presented below. Specimens that were severely bent or distorted, and those whose pectoral fins obscured their operculum, were excluded from the analysis. Landmarks placed on the left-lateral photos were used to build a Tps file using tpsUtil version 1.79 (Rohlf, 2019).

To quantify two-dimensional body shape, we digitized 27 anatomical landmarks on each of the left-lateral photographs using tpsDIG2w32, version 2.31 (Rohlf, 2018). The landmark selection was based on previously established landmarks (Krisjansson 2005, Taugbol 2014). All landmarks were visible from the lateral side of the fish (Figure 1). The number of dorsal spines varies among individuals and populations from four to six (Nelson 1969). For this reason, instead of recording the location of each dorsal spine, the location of the first and last dorsal spine were used as landmarks. In case the dorsal spine landmarks are, in fact, not homologous, we also analyzed whole-body shape data without the posterior dorsal spine landmark and without any dorsal spine landmarks. Regardless, for samples collected in 2019, it was only possible to place landmarks on the head (see Figure 1) because the posterior portion of each fish (i.e. posterior to the pelvic girdle) was destroyed for use in the stable isotope analysis. Hence, whole-body morphometric analysis used only samples collected in 2017, whereas analysis of head morphology used samples from 2017 and 2019.

**Figure 1.**
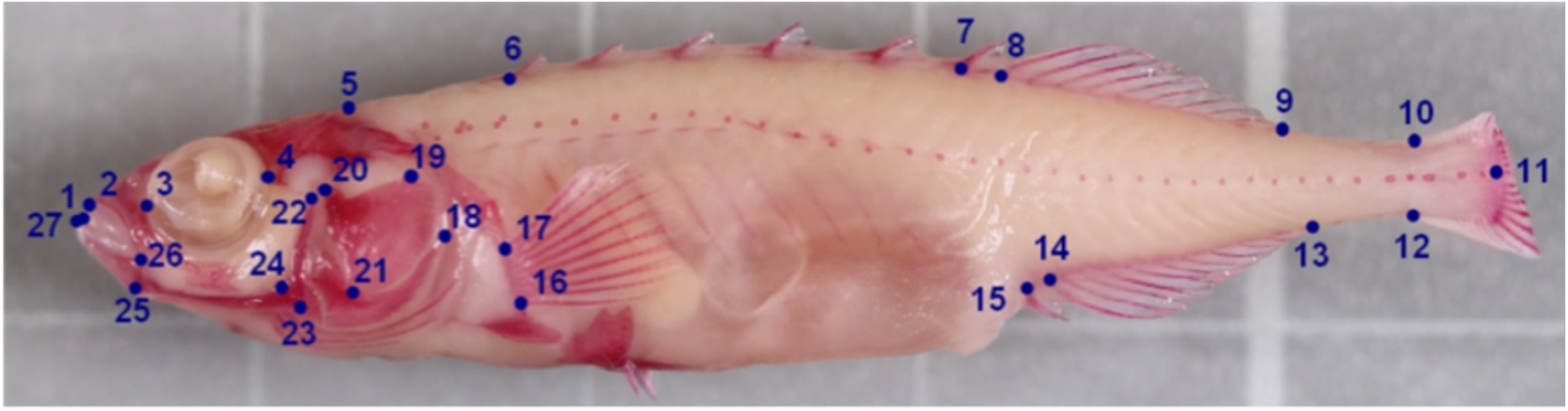
Position of 27 morphological landmark locations on a brook stickleback specimen (n = 105). 1) Anterior ventral tip of upper lip, 2) Anterior dorsal tip of upper lip, 3) Anterior border or the eye, 3) Posterior border of the eye, 4) Posterior dorsal tip of skull, 5) Base of first dorsal spine, 6) Base of last dorsal spine, 7) Anterior junction of the dorsal fin, 9) Posterior junction of the dorsal fin, 10) Dorsal insertion of the caudal fin, 11) Posterior end of the hypural plate at the midline, 12) Anterior insertion of the caudal fin, 13) Posterior junction of the anal fin, 14) Anterior junction of the anal fin, 15) Base of anal spine, 16) Anterior tip of the pectoral fin, 17) Dorsal tip of the pectoral fin, 18) Dorsal tip of the gill cover, 19) Posterior dorsal tip of operculum, 20) Anterior dorsal tip of operculum, 21) Ventral tip of operculum, 22) Dorsal tip of preoperculum, 23) Posterior angular tip of preoperculum, 24) Posterior jaw, 25) Anterior tip ventral jaw, 26) Posterior tip of lips, 27) Anterior tip of lower lip. Landmarks 7 through 15 were unavailable for samples collected in 2019.

### Stable Isotope Analysis

Stickleback, zooplankton, and benthic macroinvertebrate tissues collected in 2019 were thawed then rinsed with distilled H_2_O to clean off any lake-debris or mud. We then dried each sample at 65°C for 48 hrs in an incubator oven, ground it into a powder in liquid nitrogen using a mortar and pestle, then packed the ground tissue into 4 × 6 mm tin capsules for isotope analysis. Stable isotope analysis was performed on the packed capsules using a Carbon and Nitrogen Ratio Mass Spectrophotometer at the University of Calgary Geosciences Isotope Analysis Laboratory. Stable isotope ratios are expressed as a delta notation (δ) which is defined as the parts per million (‰) difference from a universal standard (Zanden *et al*. 1999). The standard material for δ13C is Pee Dee belemnite (PDB) limestone, and for δ15N it is atmospheric nitrogen (both ‰ values arbitrarily set at 0 ‰; Zanden *et al*. 1999, Ben-David and Flaherty 2012). To assess accuracy and repeatability of our isotope ratio measurements, we recorded triplicate or duplicate isotope measurements for 22.5% of samples (including all plankton samples).

### Statistical Analysis

To evaluate the hypothesis that pelvic phenotype is associated with other morphological changes, we assessed shape variation between pelvic phenotypes using the GEOMORPH package in R (Adams and Collyer 2020). We first performed a Generalized Procrustes analysis to estimate a scaling factor that compensated for the natural variation in fish size and applied this to all samples. This assured that all landmarks from the samples were placed on comparable locations and avoided wide dispersion of landmark coordinates. We used the procD.lm function to conduct Procrustes ANOVA (with type III sums of squares to allow for unbalanced data – see Table1), which uses a linear model to evaluate the morphological variation attributable to the following factors: lake, sex, year, pelvic phenotype, and body size. We expected that the effect of pelvic phenotype on morphology might be different between lakes or between sexes, and that the effect may be influenced by allometry (Aguirre et al. 2008, Reimchen et al. 2016). So, we included two-way interactions in our linear model. The probability of the observed effect for each factor was evaluated by comparison to a null distribution generated by 10,000 resampling permutations. We used a reverse stepwise approach for model selection, starting with the full model (main effects and two-way interactions) and removing any body size (i.e. allometric) interaction terms that were not significant.

We performed a visual evaluation of variation in isotopic signatures among fish, plankton, and benthic macroinvertebrate samples using a bi-plot of δ15N and δ13C signatures. To evaluate the hypothesis that the spined and unspined brook stickleback pelvic phenotypes use different habitats and forage on different food sources, we tested the association of sex, lake, pelvic phenotype, and fish size with δ15N and δ13C signatures using generalized linear models with gaussian error distributions. We expected that the association between pelvic phenotype and isotope signatures might be different between lakes or between sexes, and that the association may be influenced by allometry (Reimchen et al. 2016). So, as with the analysis of morphological variation, we included two-way interactions in our model. We set contrasts among factors using the contr.sum function, and we used the CAR package (Fox and Weisberg 2019) to generate an ANOVA table (with type III sums of squares to allow for unbalanced data – see Table1) to evaluate the variation in δ15N and δ13C signatures attributable to each factor and their interactions terms. We used a reverse stepwise approach for model selection, starting with the full model (main effects and two-way interactions) and removing any body size (i.e. allometric) interaction terms that were not significant. In addition, to avoid over-fitting our model (with a sample size of only 105 brook stickleback δ15N and δ13C isotope signatures), we further reduced the number of interaction terms in our analyses of stable isotope variation by removing any two-factor interactions that were not significant.

## Results

A complete summary of all samples analyzed in this study, categorized by year, lake, habitat, sex, and pelvic phenotype, is presented in Table 1. We caught no fish in the limnetic zone in Shunda Lake. The limnetic-caught fish from Muir Lake were significantly smaller than littoral-caught fish (two-sided *t* = −2.618, df = 53.437, p = 0.01148), and we only caught three female fish in the Muir Lake limnetic zone – none of which were spined. We excluded limnetic-caught fish from subsequent statistical analyses.

**Table 1.** Summary of samples analyzed in this study. Morphological analyses focused on samples from 2017, whereas analysis of stable isotopes only included samples from 2019. For all analyses described in the text, the vestigial and absent pelvic phenotypes were combined into a single “reduced” phenotypic category. All statistical analyses used type III sums of squares to account to unbalanced sampling.

|  | Spined | Pelvic Phenotype<br>Vestigial | Absent |
| --- | --- | --- | --- |
| 2017 |  |  |  |
| Muir Lake |  |  |  |
| Littoral |  |  |  |
| female | 20 | 1 | 24 |
| male | 24 | 1 | 18 |
| Shunda Lake |  |  |  |
| Littoral |  |  |  |
| female | 20 | 24 | 9 |
| male | 29 | 6 | 3 |
| 2019 |  |  |  |
| Muir Lake |  |  |  |
| Littoral |  |  |  |
| female | 3 | 0 | 4 |
| male | 16 | 0 | 15 |
| Limnetic |  |  |  |
| female | 0 | 0 | 3 |
| male | 6 | 1 | 10 |
| Shunda Lake |  |  |  |
| Littoral |  |  |  |
| female | 14 | 11 | 4 |
| male | 18 | 9 | 2 |

### Morphology

We organized our analyses of morphological variation based on which landmarks were available among samples from different years: one analysis involving sixteen head-only landmarks that were present in all samples (from 2017 and 2019: number of observations = 295), and one analysis involving all twenty-seven landmarks, some of which (i.e. posterior to the pelvic girdle) were not present in 2019 samples because of destructive sampling for stable isotope analysis (number of observations = 179). For simplicity, Figure 2 shows only the whole-body morphological associations. Brook stickleback head-body morphology was significantly associated with fish size and differed significantly between sexes (2017 whole-body: Table 2; 2017-2019 heads only: Table 3). In addition, head morphology varied significantly among years (Table 3). Females had more elongated abdominal regions, narrower bodies, and shorter heads, which is a pattern observed in threespine stickleback as well (Aguirre et al. 2008). The association between pelvic phenotype and head-body morphology was different in each lake. In Muir Lake, pelvic reduction was most-noticeably associated with a deeper body, larger head, and anterior-shifted pectoral fins, whereas in Shunda Lake, pelvic reduction was associated with a compression of the ventral portion of the head (including a less up-curved mouth), posterior-shifted pectoral fins, and a longer, narrower body (Figure 2). These patterns are consistent with our hypothesis that brook stickleback morphology is affected by pelvic phenotype, and these results are consistent with previous observations that morphological divergence is non-parallel (Kaeuffer et al. 2012).

**Table 2.** ANOVA table for the linear model fitted to whole-body 2D morphology for brook stickleback collected in 2017.

|  | df | Type III SS | F | p |
| --- | --- | --- | --- | --- |
| Log(length) | 1 | 0.004158 | 4.4343 | **0.0008** |
| Lake | 1 | 0.008727 | 9.3065 | **< 0.0001** |
| Sex | 1 | 0.014800 | 15.7835 | **< 0.0001** |
| Pelvic phenotype | 1 | 0.002982 | 3.1796 | **0.0086** |
| Lake * Sex | 1 | 0.001971 | 2.1021 | 0.0532 |
| Lake * Pelvic phenotype | 1 | 0.003046 | 3.2482 | **0.0075** |
| Sex * Pelvic phenotype | 1 | 0.000754 | 0.8043 | 0.5432 |
| Residuals | 171 | 0.160345 |  |  |

**Table 3.**
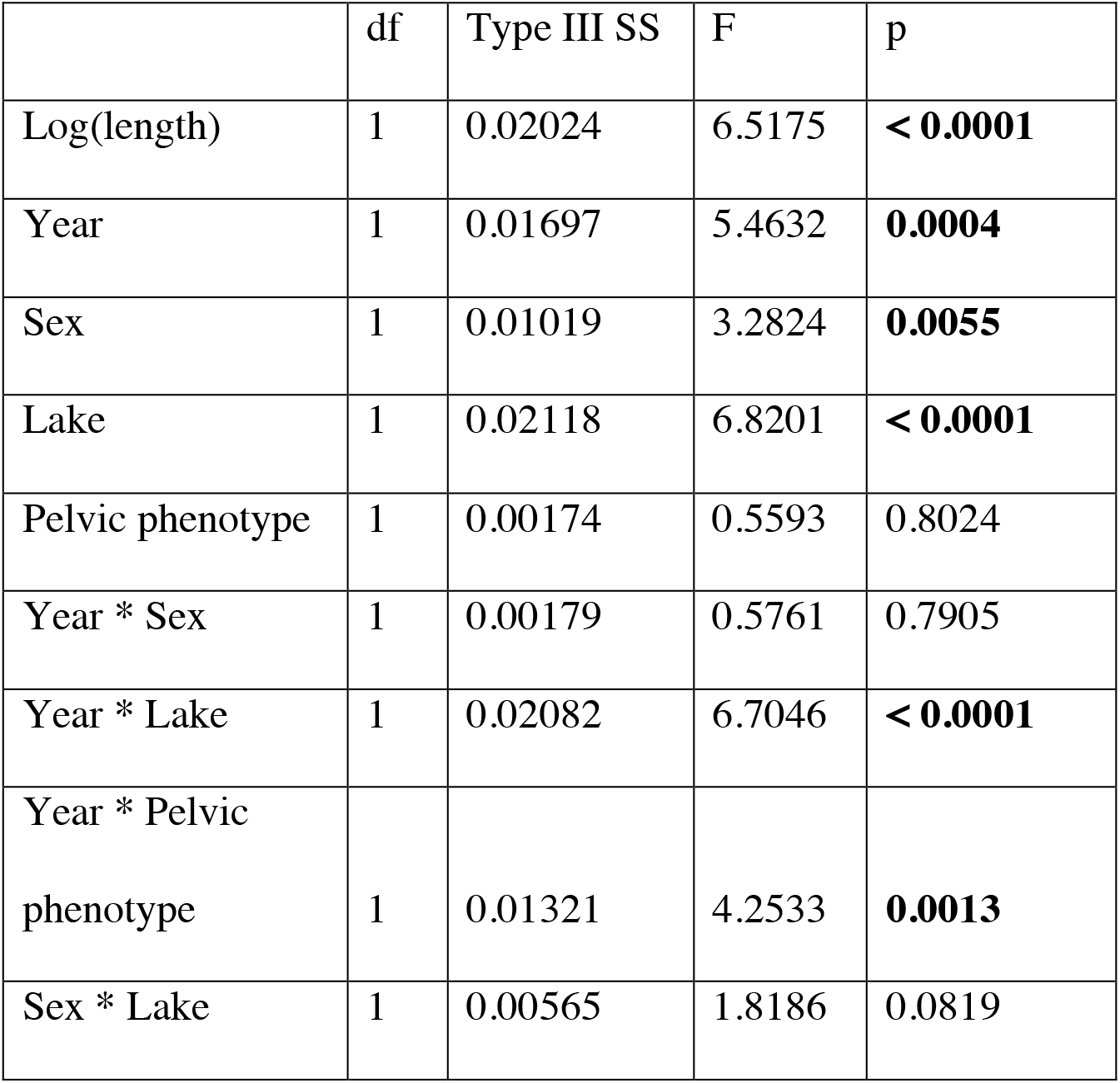
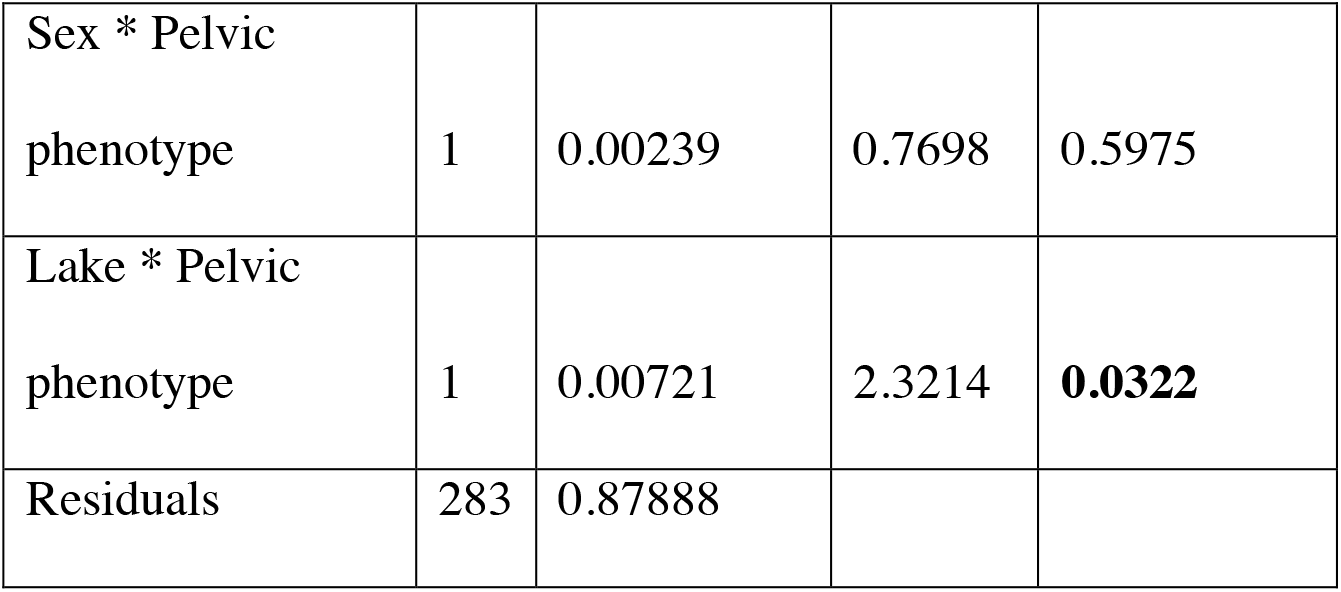
ANOVA table for the linear model fitted to head-only 2D morphology for brook stickleback collected in 2017 and 2019.

**Figure 2.**
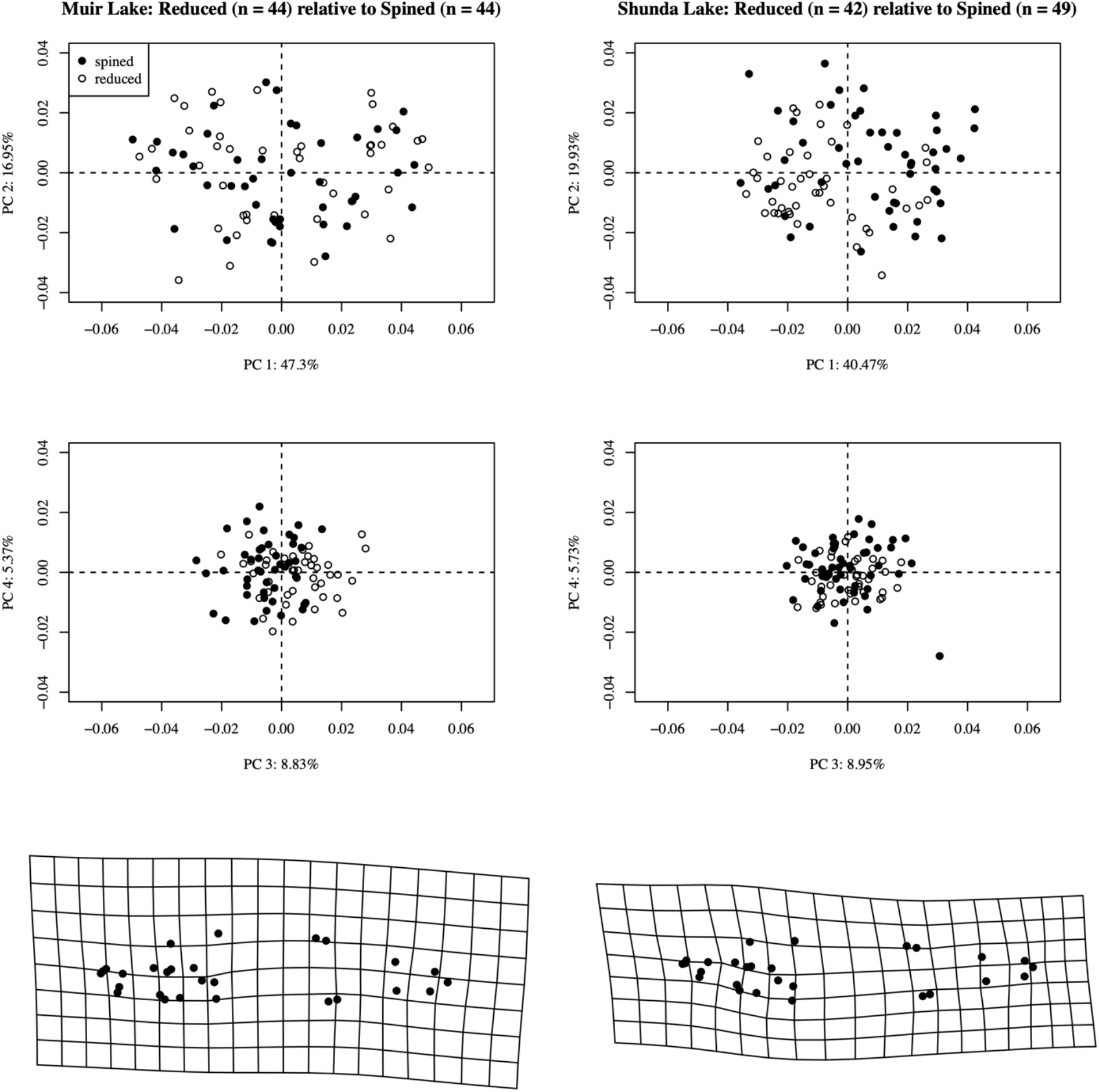
The effect of pelvic phenotype on body morphology in brook stickleback from Muir Lake (left column) and Shunda Lake (right column). The top two plots in each column show the results of principle components analyses for each lake. Values for each individual fish for the first four principle components (PCs) are shown with the proportion variance in body morphology explained by each PC shown on the axis labels. The bottom plot in each column shows the distortion of a symmetrical (square) grid when the average shape of a brook stickleback with a reduced pelvic phenotype is superimposed on the average shape of a brook stickleback with a complete pelvic structure.

### Stable isotopes

In both lakes, brook stickleback had higher δ15N isotopic signatures than benthic invertebrates (Figure 3) suggesting that, as expected, fish occupy a higher trophic niche relative to the macroinvertebrates. In Shunda Lake, the plankton had the lowest δ15N signature (Figure 3) suggesting the expected hierarchy in trophic position, with fish at the top, plankton at the bottom, and macroinvertebrates in the middle. In Muir Lake, the δ15N isotopic signature was not substantially lower than the average for macroinvertebrates, but the δ13C signature for plankton in Muir Lake was substantially higher than any of the other samples. The higher-than-expected δ13C signature and higher-than-expected δ15N signature for plankton in Muir Lake is an unexpected result, and further investigation is needed for a satisfactory explanation.

**Figure 3.**
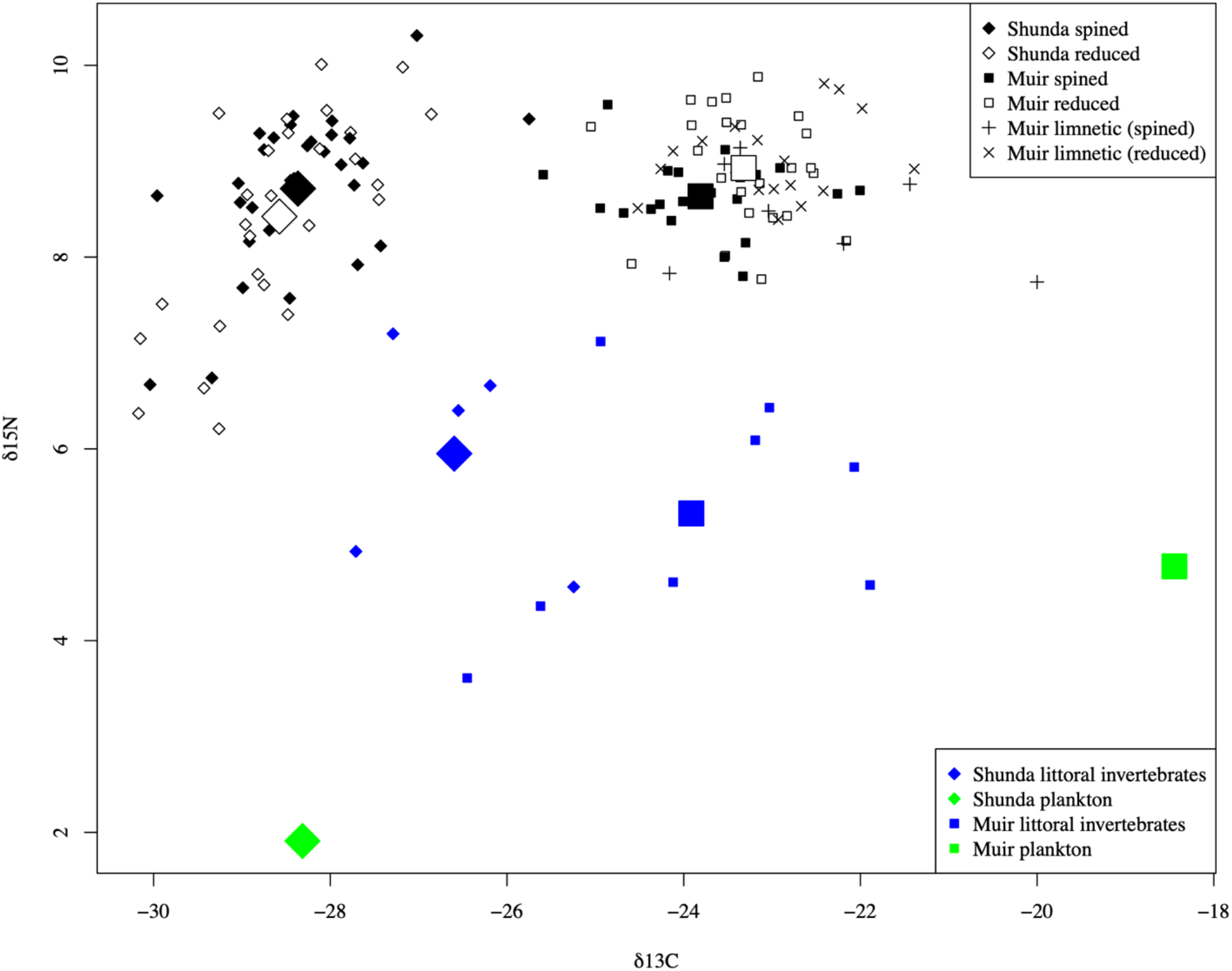
Carbon and nitrogen stable isotope signatures for spined and unspined brook stickleback, zooplankton, and benthic littoral macroinvertebrates from Muir Lake and Shunda lake. Macroinvertebrate taxa represented in the plot include caddisfly larvae in the family Limnephilidae, unidentified caddisfly larvae, amphipods, leaches, dragonfly larvae in the family Gomphidae, unidentified dragonfly larvae, unidentified snails, and unidentified damselfly larvae. Also shown (with + and x symbols) are limnetic-caught brook stickleback from Muir Lake. All fish were caught in the littoral zone unless otherwise indicated. The larger symbols represent mean values for each subgroup (excluding limnetic-caught), whereas the smaller symbols represent values for individuals.

Stickleback δ13C isotope signatures were significantly associated with fish size (Table 4), but none of the interactions terms with fish size were significant. This suggests that the effect of size on δ13C signature is the same regardless of pelvic phenotype, sex, or lake. The association between pelvic phenotype and δ13C signature was different in each lake, and Muir Lake had a much higher δ13C signature than Shunda Lake (Table 4, Figure 3). In Muir Lake, pelvic reduction was associated with a higher δ13C signature, whereas in Shunda Lake, pelvic reduction was associated with a lower δ13C signature (Figure 4). In both lakes, however, the shift in brook stickleback δ13C signature associated with pelvic reduction was towards the planktonic δ13C signature (Figure 3). In Muir Lake, limnetic-caught fish tended to have a higher δ13C signature than littoral-caught fish (Figure 3). These patterns are consistent with our hypothesis that brook stickleback with different pelvic phenotypes forage in different habitats, but these patterns are inconsistent with our prediction that, as in threespine stickleback, pelvic reduction would be associated with littoral or benthic habitats. In fact, contrary to our prediction, these results suggest that pelvic reduction is associated with limnetic (or planktonic) feeding in both lakes.

**Table 4.** ANOVA table for the generalized linear model fitted to δ13C isotopic signature for brook stickleback collected in 2019.

|  | df | Type III SS | F | p |
| --- | --- | --- | --- | --- |
| Lake | 1 | 527.5 | 848.4586 | **< 0.0001** |
| Pelvic phenotype | 1 | 0.29 | 0.4613 | 0.49873 |
| Sex | 1 | 0.64 | 1.0329 | 0.31214 |
| Length | 1 | 4.69 | 7.5448 | **0.00724** |
| Lake * pelvic phenotype | 1 | 2.97 | 4.7848 | **0.03125** |
| Residuals | 92 | 57.2 |  |  |

**Figure 4.**
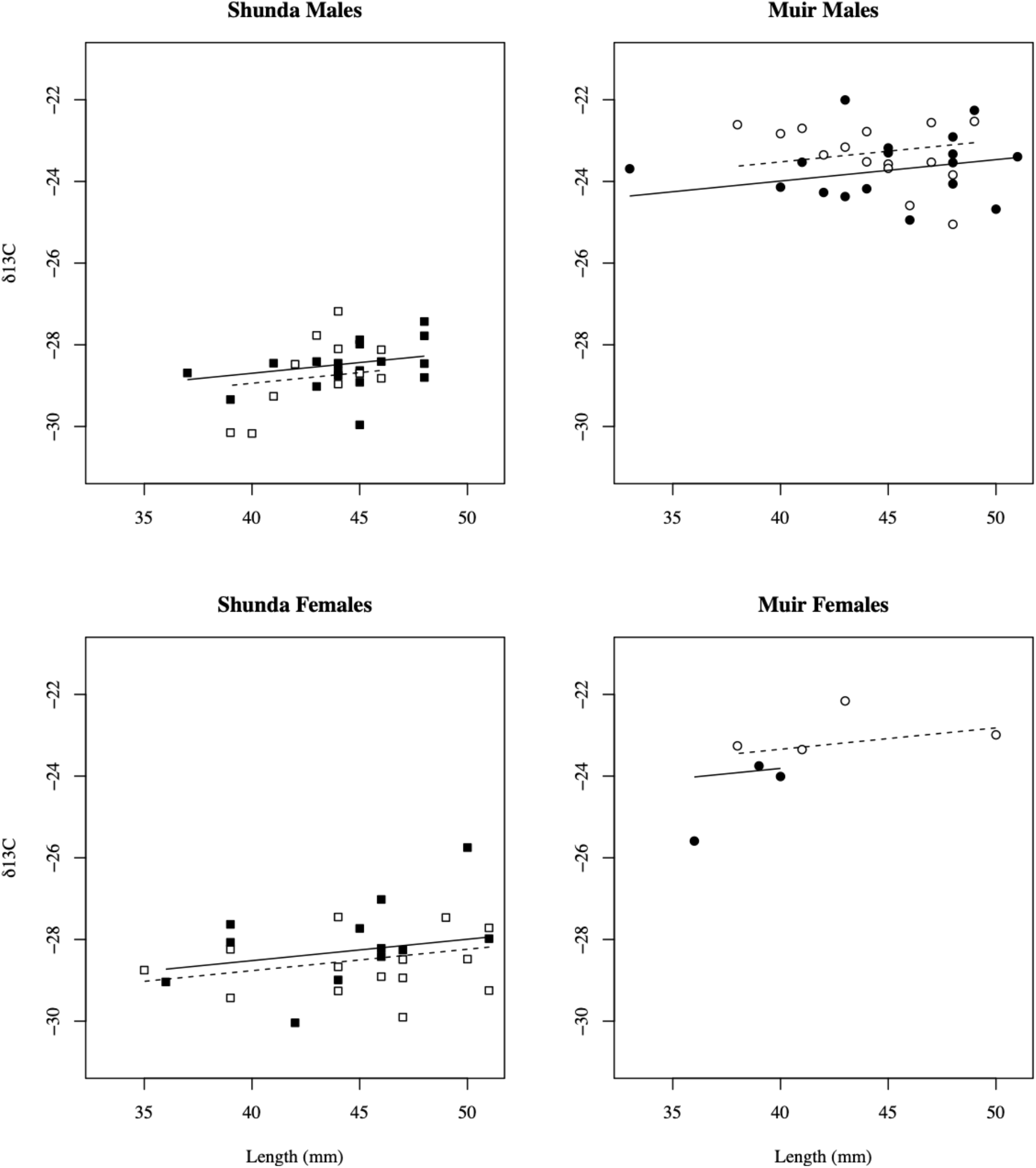
Summary of the effects of sex, lake, pelvic phenotype, and fish size on δ13C isotope signature in brook stickleback. As in Figure 3, closed symbols represent spined fish and open symbols represent fish with reduced pelvic phenotypes. Lines show the linear relationships inferred from the model described in the text and in Table 4 (solid = spined, dashed = reduced).

The effect of size on stickleback δ15N isotopic signature was dependent on lake (Table 5). In Muir Lake, larger fish had a lower δ15N isotopic signature, whereas larger fish had a higher δ15N isotopic signature in Shunda Lake (Figure 5). The effect of pelvic phenotype on stickleback δ15N isotopic signature was dependent on sex. In males, pelvic reduction is associated with a higher δ15N signature, whereas pelvic reduction is associated with a lower δ15N signature in females (Figure 5). These patterns are consistent with our hypothesis that brook stickleback pelvic morphs forage in different habitats.

**Table 5.**
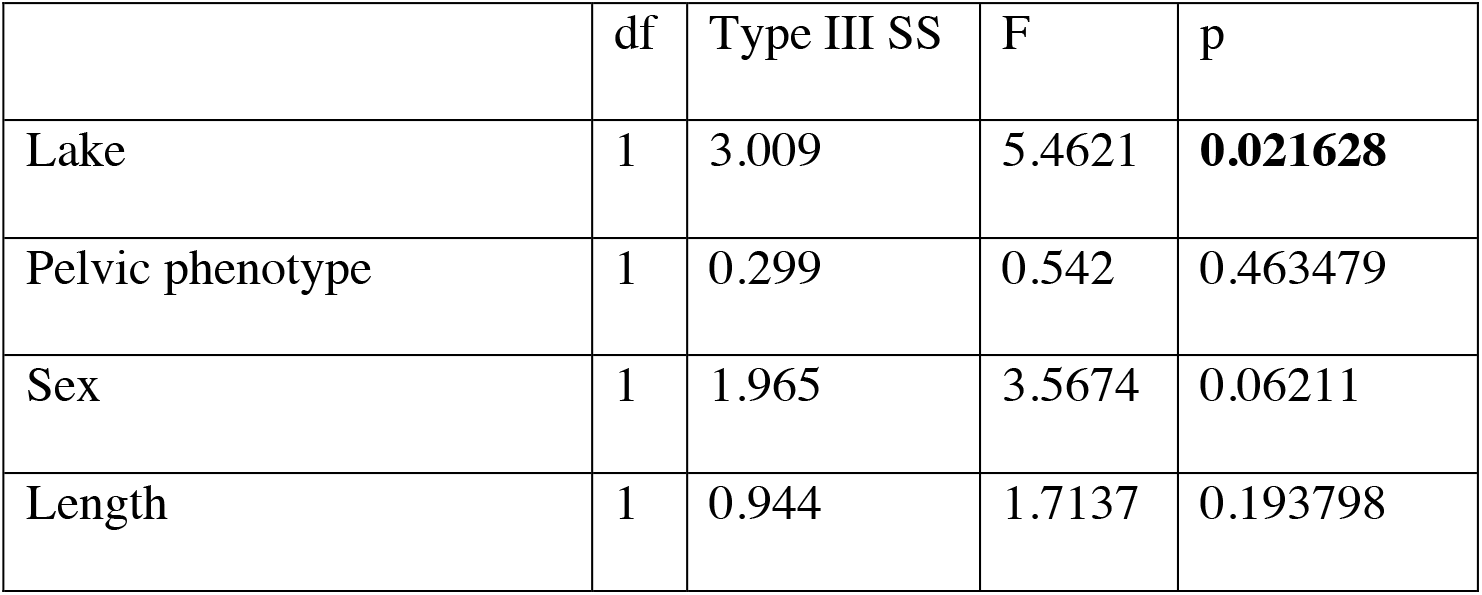
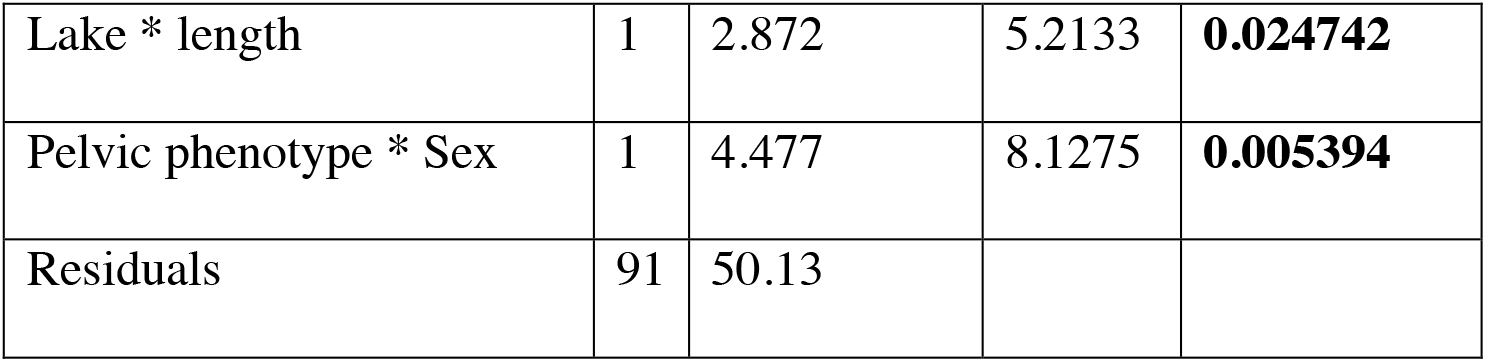
ANOVA table for the generalized linear model fitted to δ15N isotopic signature for brook stickleback collected in 2019.

|  | df | Type III SS | F | p |
| --- | --- | --- | --- | --- |
| Lake | 1 | 3.009 | 5.4621 | **0.021628** |
| Pelvic phenotype | 1 | 0.299 | 0.542 | 0.463479 |
| Sex | 1 | 1.965 | 3.5674 | 0.06211 |
| Length | 1 | 0.944 | 1.7137 | 0.193798 |
| Lake * length | 1 | 2.872 | 5.2133 | **0.024742** |
| Pelvic phenotype * Sex | 1 | 4.477 | 8.1275 | **0.005394** |
| Residuals | 91 | 50.13 |  |  |

**Figure 5.**
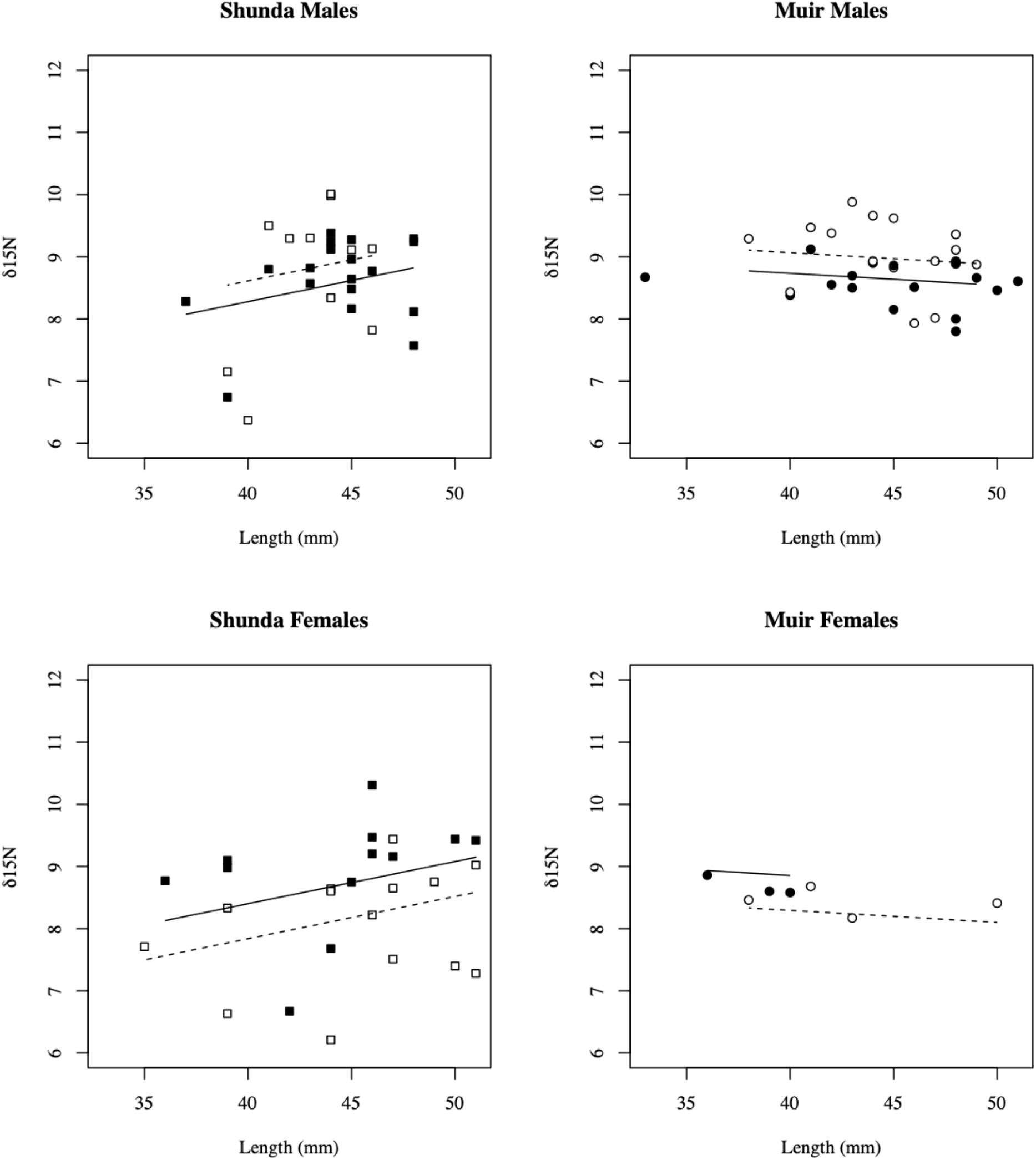
Summary of effects of sex, lake, pelvic phenotype, and fish size on δ15N isotope signature in brook stickleback. As in Figure 3, closed symbols represent spined fish and open symbols represent fish with reduced pelvic phenotypes. Lines show the linear relationships inferred from the model described in the text and in Table 5 (solid = spined, dashed = reduced).

## Discussion

The hypothesis that pelvic spine polymorphism in brook stickleback is associated with divergence in habitat use was supported by our results. Based on carbon isotope signatures, brook stickleback with pelvic reduction (i.e. with either no pelvic structure or a vestigial pelvic structure) likely feed on planktonic as opposed to benthic macroinvertebrate food sources and, therefore, likely use more limnetic as opposed to littoral or benthic habitat. This result is, however, contrary to our prediction based on the well-established association between pelvic reduction and benthic habitats in threespine stickleback (Reimchen 1980, McPhail 1992). There were also significant changes in head and body morphology associated with pelvic reduction, but, as expected (Kaeuffer et al. 2012), the specific nature of this morphological difference was lake dependent, and the magnitude of the morphological difference between pelvic phenotypes was small. In fact, the proportion of the variation in morphology, δ13C isotope signature, or δ15N signature attributable to pelvic spine variation was small relative to individual-level variation and between-lake variation (Figures 3 and 4, Tables 2, 3, 4, and 5).

There is ample evidence supporting the role of stickleback pelvic spines in predator interactions. It is likely that balancing selection driven by multiple and varied predators is involved in maintaining the spine polymorphism, and that predation drives associations between spine morphology and diet, habitat, and body morphology. But, we do not know the mechanism via which predation and other ecological factors cause the unexpected association between pelvic reduction and a more planktonic diet (as opposed to the predicted association between pelvic reduction and a more benthic diet). It is possible that, in these systems, stocked trout (or other gape-limited predatory fish) forage in littoral zones, thereby selecting for spined stickleback in these habitats. Stocked rainbow trout have been observed ambushing minnows from under floating docks in the littoral zones of other nearby lakes (J. Mee personal observation).

We found that, in unspined fish only, males had significantly higher δ15N signatures than females. This suggests that unspined females occupy a lower trophic position than unspined males, which may indicate than unspined females have a more planktivorous diet, whereas unspined males have a more benthic diet. This is consistent with patterns observed in male threespine stickleback, which are more benthic-associated than female threespine stickleback as a consequence of their breeding behaviour (Spoljaric and Reimchen 2008, Aguirre and Akinpelu 2010, McGee and Wainwright 2013). Male threespine stickleback migrate from the limnetic to the littoral habitats to build nests, spawn, and defend their eggs. Brook stickleback also build nests and defend their eggs, but, whereas threespine stickleback build their nests on the substrate, brook stickleback males build nests on vertical rocky surfaces or vegetation (Reisman and Cade 1967, McLennan 1995). Hence, it is not clear that the breeding behaviour of brook stickleback males should cause them to be more benthic-associated than females. Also, if the difference in δ15N signature between unspined males and females is due to male breeding behaviour, it is not clear why spined males do not have a higher δ15N signature than spined females unless the breeding behaviour of spined and unspined males is different. It seems likely that differences between brook stickleback and threespine stickleback life history and reproductive behaviour obscure any simple comparison between the species regarding the interactions among diet, habitat, and morphology.

Morphological variation among threespine stickleback populations is associated with habitat specialization (Reimchen et al. 1985, Webster 2011). Even subtle changes in body morphology can be associated with fitness parameters like foraging patterns, body condition, and growth rate (Webster et al. 2011). Different selection pressures in different habitats may favour different morphological traits (Reimchen et al. 1985). If traits that are well-adapted for a certain environment are ill-suited for another, there may be fitness trade-offs among habitats (Webster et al. 2011). The morphological variation among pelvic phenotypes may be due to differing water depth, water chemistry (e.g. pH), predation risk, and parasite prevalence in different habitats (Reimchen et al. 1985, Webster 2011). There is, therefore, good reason to assume that differences in habitat use and diet between spined and unspined brook stickleback cause the observed morphological differences between morphs in the present study. However, it is not clear if the morphological differences between spined and unspined brook stickleback constitute a plastic response or a heritable response to divergent habitat use. Additionally, if the morphological divergence is heritable, we do not know whether it has functional significance driven by different selective regimes in different habitats, or if it results from pleiotropic effects of the genes underlying the pelvic divergence.

There was an obvious difference in δ13C signature between the population in Muir Lake and the population in Shunda Lake. This could be the result of different diet preferences between populations (Jardine *et al*. 2003, Eloranta et al. 2010), or it may reflect chemical differences in the environment (e.g. weather, sediment, or human impact, pH). Muir Lake is in Alberta’s Central Parkland natural region, which is a prairie landscape, and its riparian area is dominated by marsh vegetation such as cattails (*Typha* spp.) and sedges (*Carex* spp.), and likely has relatively high pH (AWA 2020). Shunda Lake is in the Upper Foothills natural region, which is mountainous, and its riparian area is characterized by conifer stands and understory shrubs, and likely has relatively low pH (AWA 2020, Ross and Kyba 2015). This difference between lakes may be a reason for the difference in isotopic signatures among the plankton and invertebrate samples in these two lakes and may represent an ecological basis for the differences in morphology and isotopic signature between these two brook stickleback populations.

The isotopic signatures of brook stickleback, benthic invertebrates, and zooplankton relative to one another (Figure 3) were generally consistent with the linear ascending relationship of δ15N in food-web and trophic position studies of freshwater fish and their food sources (Post 2002). The zooplankton signature from Muir Lake did not, however, fit neatly within this paradigm. We lack any evidence to support an explanation for this unexpected isotopic signature in Muir Lake, but we can offer some conjecture. It is possible that Muir lake, at the time of sampling, was dominated by a single species or relatively few species of zooplankton (n.b. we did not identify zooplankton to species in our study). The expectation that zooplankton should have a lower δ15N and a higher δ13C isotopic signature than their consumers (Post 2002) assumes an average isotopic signature among a community of zooplankton. If our sample constituted only a few species (or even a single species), it may have been a species with a particularly 15N-enriched and 13C-enriched isotopic signature. Further sampling and analysis of isotopic signatures in this lake would be required to explain this result.

The observation of significant differences in morphology and stable isotope signature between lakes suggests another future avenue of inquiry. If differences between lakes are heritable, there may be implications related to parallelism in the evolution of pelvic spine reduction. It is unknown whether the genetic basis of pelvic spine polymorphism is the same in all brook stickleback populations. If differing body morphologies between lakes are associated with different genetic bases for pelvic reduction in different lakes (e.g. via pleiotropy), there may be less phenotypic and genetic parallelism in this system than previously assumed. Investigations of the heritability and genetic bases of polymorphism in these lakes are ongoing.

## Acknowledgments

Three anonymous reviewers provided insightful and constructive comments on a previous version of this manuscript. Jim Reist provided feedback on an earlier draft of this manuscript. Arianne Albert and Mike Rennie provided feedback on statistical methods. Mike Rennie and Olivier Morrissette provided input on the collection and analysis of stable isotope data. We would like to thank Melanie Rathburn, Dorothy Hill, and Matthew Swallow for their guidance, encouragement, and constructive feedback throughout the project. Special thanks to everyone who participated in sample collection: Alex Farmer, Carolyn Ly, Alyce Straub, Bryce Carson, Moroni Lopez Vasquez, and Kaylee Olszewski. Steven Taylor at the University of Calgary Geosciences Isotope Analysis Laboratory trained EF in stable isotope sample preparation and analysis, and assisted with the isotope data collection. This work was supported by a grant from the Institute for Environmental Sustainability at Mount Royal University, and by a Research Grant from the Office of Research, Scholarship, and Community Engagement. An NSERC Discovery Grant awarded to JAM supported this work, and KW was supported by an NSERC URSA.

## Shared Data

All data and analysis scripts have been uploaded to the Dryad Data repository: https://datadryad.org/stash/share/Vho8zW5Aqhj5yACQtLSKf0Dm9BWcpGXIHLIVJshTeQ0

